# Conformational entropy limits the transition from nucleation to elongation in amyloid aggregation

**DOI:** 10.1101/2020.06.22.165423

**Authors:** Tien M. Phan, Jeremy D. Schmit

**Affiliations:** Department of Physics, Kansas State University, Manhattan, KS 66506, USA

## Abstract

The formation of *β*-sheet rich amyloid fibrils in Alzheimer’s disease and other neurodegenerative disorders is limited by a slow nucleation event. To understand the initial formation of *β*-sheets from disordered peptides, we used all-atom simulations to parameterize a lattice model that treats each amino acid as a binary variable with *β* and non-*β* states. We show that translational and conformational entropy give the nascent *β*-sheet an anisotropic surface tension which can be used to describe the nucleus with two-dimensional Classical Nucleation Theory. Since translational entropy depends on concentration, the aspect ratio of the critical *β*-sheet changes with protein concentration. Our model explains the transition from the nucleation phase to elongation as the point where the *β*-sheet core becomes large enough to overcome the conformational entropy cost to straighten the terminal molecule. At this point the *β*-strands in the nucleus spontaneously elongate, which results in a larger binding surface to capture new molecules. These results suggest that nucleation is relatively insensitive to sequence differences in co-aggregation experiments because the nucleus only involves a small portion of the peptide.

**SIGNIFICANCE:** The conversion of soluble proteins to amyloid aggregates is associated with many neurodegenerative diseases. Experiments have shown that this conversion occurs by a slow nucleation step followed by rapid growth. This work identifies the principle contributions to the free energy barrier that separates these two stages. It also shows how factors like protein concentration, sidechain interactions, and interactions with the environment can modify the barrier and affect nucleation times.

## INTRODUCTION

The assembly of proteins into amyloid fibrils causes numerous neurodegenerative diseases, such as Alzheimer’s disease, Parkinson’s disease, and prion disorders (1). *In vitro* experiments show that the conversion of soluble peptides to the fibril state is limited by nucleation events, indicative of a free energy barrier to initiate the fibril state. The linear morphology of amyloid fibrils suggests that the nucleation barrier must be substantially different than other nucleating systems. In the case of isotropic particle condensation, the nucleation barrier arises because particles at the periphery have sacrificed the translational entropy of the dilute phase, but only form a fraction of the favorable interactions available to interior particles (2). This interaction deficit, usually described as a surface tension, becomes a smaller fraction of the free energy as the cluster grows larger.

Surface tension does not limit 1D assemblies because the surface energy does not depend on the cluster size (3). Early attempts to explain amyloid nucleation identified *β*-sheet layering as a second assembly dimension (4–9). An alternative mechanism that can result in nucleation behavior in 1D systems is a conformational conversion to the *β*-sheet state (10). This mechanism has been implicitly included in coarse-grained simulations by modeling the molecules with two states; a high energy but strongly attractive fibrillar state, and a low energy state with weaker interactions (11–13). Due to the large energy difference between these states, it was found that non-fibrillar oligomers were necessary to overcome the energy barrier to initiate a fibril. In this paper we remove the two-state assumption by allowing the molecule to partially convert to the fibril state. This partial conversion enables pathways with a lower barrier to conformational conversion. We find that the nucleation of a *β*-sheet, and hence an amyloid fibril, is limited by two contributions to the free energy barrier, one arising from the translational entropy cost to recruit molecules from solution, and one arising from the conformational entropy cost to form *β*-strands. These contributions can be modeled using Classical Nucleation Theory (CNT) with an anisotropic surface tension. The distinct entropic contributions contributing to the surface tension in different directions means that the *size and shape* of the critical *β*-sheet is sensitive to factors like the amino acid sequence and protein concentration. We also find that there is a critical fibril length at which the conformational entropy barrier vanishes. At this point the *β*-strands in the cluster will spontaneously elongate. These longer *β*-strands provide a stronger binding surface for incoming molecules, which we attribute to the end of the nucleation phase and the beginning of the elongation phase.

## METHOD

We employ a dual resolution model to simulate the conversion of peptide strands from the disordered state to a *β*-sheet. Our approach follows from recent work in which fibril elongation was modeled by treating each amino acid as a binary variable that can be in either *β* or non-*β* states (14, 15). We use all-atom molecular dynamics (MD) on a hexamer and dimer of polyglutamine, motivated by the aggregation-prone region of huntingtin protein (16–23), to measure kinetic transitions between the *β* and non-*β* states in the post- and pre-nucleation scenarios, respectively (Fig. 1).

**Figure 1:**
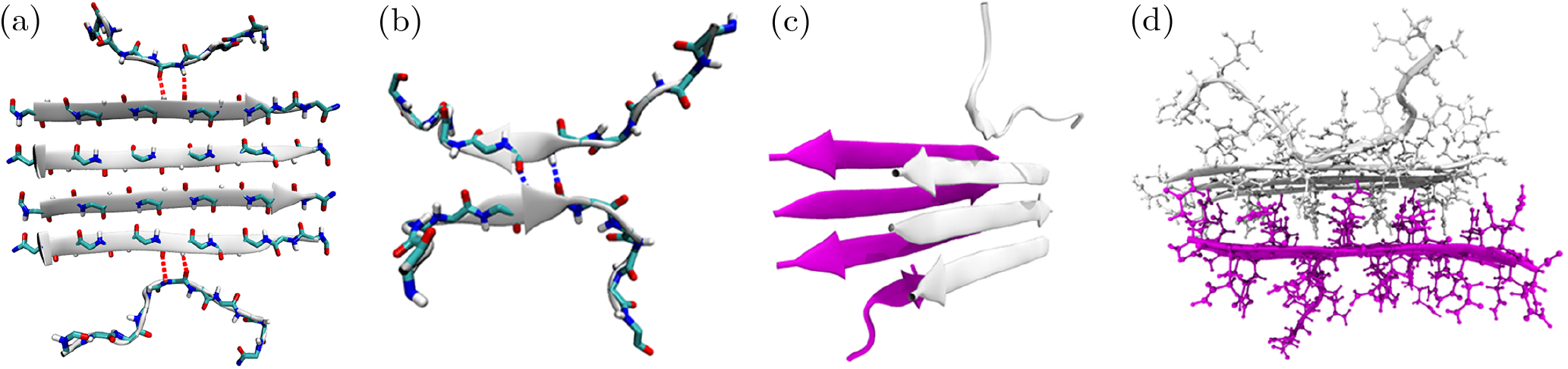
MD snapshots from the sampling of the conversion between *β* and non-*β* states. (a) Kinetic parameters for post-nucleation (“strong”) bonds are sampled from the terminal molecules on an established cluster (side chains are not shown for clarity). (b) Pre-nucleation (“weak”) bonds are sampled using a dimer that is harmonically restrained at the central amino acids. (c) Ribbon representation of the *β*-sheet bilayer simulation showing the stacking of the two sheets. The six core molecules are restrained in *β* conformation. The top molecule in the white sheet and the bottom molecule in the magenta sheet are free to sample *β* and non-*β* states. This side view shows the staggered alignment of *β*-strands in which the white strands are positioned between the magenta strands in the opposite sheet. This staggering ensures that each molecule addition results in an equivalent set of H-bond and sidechain interactions. (d) Atomistic view of the bilayer showing the interdigitation of side chains between the two *β*-sheets.

### Molecular dynamics simulation details

To simulate post-nucleation bonds in the all-atom model we used a *β*-sheet consisting of six monomers. The backbone atoms of the four internal molecules were harmonically restrained to mimic the rigidity of a mature fibril, along with the central amino acid of the terminal molecules, using a force constant of 10 kcal/mol/Å^2^. The system was heated to 900 K for 100 ps generating an ensemble of 10 initial states. We then monitored H-bond transitions around the anchored amino acid (Fig. 1a). Pre-nucleation bonds were assessed using an anti-parallel dimer with harmonic restraints on the central amino acids (Fig. 1b). We ran 10 replicas for each starting state lasting for 100 ns each. H-bond transition rates were obtained as described in (14).

All simulations were performed with OpenMM 7.3.1 (24) using three force fields, CHARM36m (25), AMBER99SB-ILDN (26) and AMBER14SB (27), with the TIP3P (28) water model. These force fields are suggested candidates for amyloid peptide assembly based on the study of A*β*_16–22_ dimer in (29). For all simulations, long-range electrostatic interactions were treated with particle mesh Ewald (PME), with both direct-space PME and Lennard-Jones potentials making use of a 10 Å cutoff; the Lennard-Jones potential was switched to zero at the cutoff over a switch width of 1.5 Å to ensure continuity of potential and forces. PME used a relative error tolerance of 10^4^ at the cutoff to automatically select the *α* smoothing parameter, and the default algorithm in OpenMM was used to select Fourier grid spacing (which selected a grid spacing of 0.8 Å in each dimension). All bonds to hydrogen were constrained to a within a fractional error of 10^−8^ of the bond distances using CCMA (30, 31), and waters were rigidly constrained with SETTLE (32). OpenMM’s long-range analytical dispersion correction was used to avoid pressure artifacts from truncation of the Lennard-Jones potential. Simulations were run at 300 K with a Monte Carlo barostat with 1 atm external pressure and Monte Carlo update interval of 25 steps. Langevin dynamics was used with a 2 fs time steps and collision of 1 ps^−1^.

Coordinates were saved every 5 ps and the rate constants between two states, *β*-sheet and non-*β* conformations, were measured using the method in (14). A transition involves either the formation or breakage of a pair of backbone H-bonds between two residues. The distances between amide hydrogen and carbonyl oxygen pair d_H−O_ were computed using pytraj (33) and monitored for transitions. A backbone H-bond is considered broken when d_H–O_ exceeds 3.5 Å and formed when d_H–O_ is shorter than 2.5 Å. In addition to the 1 Å gap in formation and breakage cutoff distances, a five-frame (25 ps) running average of d_H–O_ and elimination of fast transition less than 300 ps were used to suppress spurious high frequency fluctuations in the detection of backbone H-bond transitions.

To approximate the free energy of the sidechain interactions with the second *β*-sheet, we repeat the MD simulation using a bilayer of 8 molecules, each of which has 10 amino acids (the pdb structure is taken from Ref. (34)). The two layers are arranged in steric-zipper orientaion in which the side chains of opposing *β*-sheets are interdigitated and one sheets is shifted upward by half of a strand relative to the other sheet (Fig. 2c,d). The central amino acids of the top and bottom molecules and the six core molecules were harmonically restrained using a force constant of 10 kcal/mol/Å^2^. The system was heated to 900 K for 100 ps generating an ensemble of 10 initial states. We ran 10 replicas for each state for 100 ns each and measured H-bond transitions around the anchored amino acid as in the monolayer case. The average time for unbound and bound states are 5272.9 ps and 25649.5 ps respectively, which yield the free energy of −1.58 *k_B_T*.

**Figure 2:**
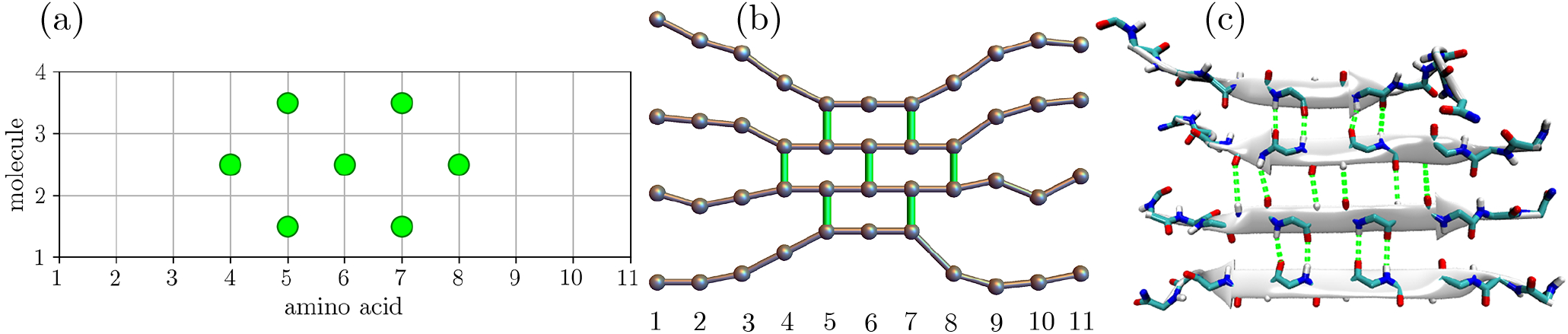
Schematic of the mapping between the lattice and all atom models shown in three representations: (a) lattice (b) ball-and-stick and (c) atomistic (side chains are hidden for clarity). The lattice has a width given by the number of amino acids per molecule and a height given by the current number of molecules in the cluster. Lattice sites (represented by vertical lines) can be occupied, representing an H-bond between connected amino acids, or empty, indicating that at least one of the adjacent amino acids is in the random coil state. The alternating direction of H-bonds (seen in the atomistic view, panel (c)) means that only every-other site can have a bond (green circles in panel (a)).

### Lattice simulation details

To observe the kinetics of nucleation we simulate the formation of *β* -sheets using a lattice model evolving via the Gillespie algorithm (35). Lattice model monomers have 11 amino acids and form anti-parallel *β*-sheets, which is more stable than parallel *β*-sheets for polyglutamine (36). Each peptide unit is modeled as a binary variable with states representing *β*-sheet and non-*β* conformations. Each amino acid can form a pair of H-bonds, which are mapped to a single bond in the lattice model (Figs. 2). Peptide units at the periphery of the *β*-core fluctuate between *β*-sheet and non-*β* states at rates measured from the all-atom model. New molecules can add to either end of the *β* -sheet at a concentration-dependent rate approximated by the Smoluchowski formula for an absorbing sphere

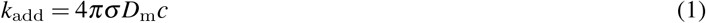

where *c* is the protein concentration, σ = 1.75 nm is the radius of the sphere, approximated by half the length of an extended monomer, and *D*_m_ is the diffusion coefficient of the monomer, 1.79 × 10^−12^m^2^/s (34).

We define the committor as the probability that a nucleus with a specified number of bonds reaches the cutoff size of 8 molecules before completely dissolving. The committor is calculated as a function of the number of molecules and the total number of H-bonds using 10,000 lattice simulations at each monomer concentration. Each simulation was initiated with a dimer with one H-bond and terminated when there were either 8 molecules in the cluster or all H-bonds were broken. The formation and breakage of H-bonds in the lattice model are described by the hexamer all-atom simulation provided one of the strands remains in the *β*-state when the bond is broken. Alternatively, if both strands are free to explore non-*β* states following the breakage of a bond we use the parameters of the dimer simulation. A new molecule can attach to either end of the *β*-sheet when it has at least two backbone H-bonds.

## RESULTS

### Simulations show that pre-nucleation bonds are much weaker than post-nucleation bonds

Figure 3 shows the probability that a peptide tail in the all-atom simulation retains the same number of H-bonds as a function of waiting time. These probabilities were fit to an exponential in which the fitting parameter is the bond breakage or formation rate. These rates vary only weakly with position or free chain length allowing us to aggregate the data for better statistics.

**Figure 3:**
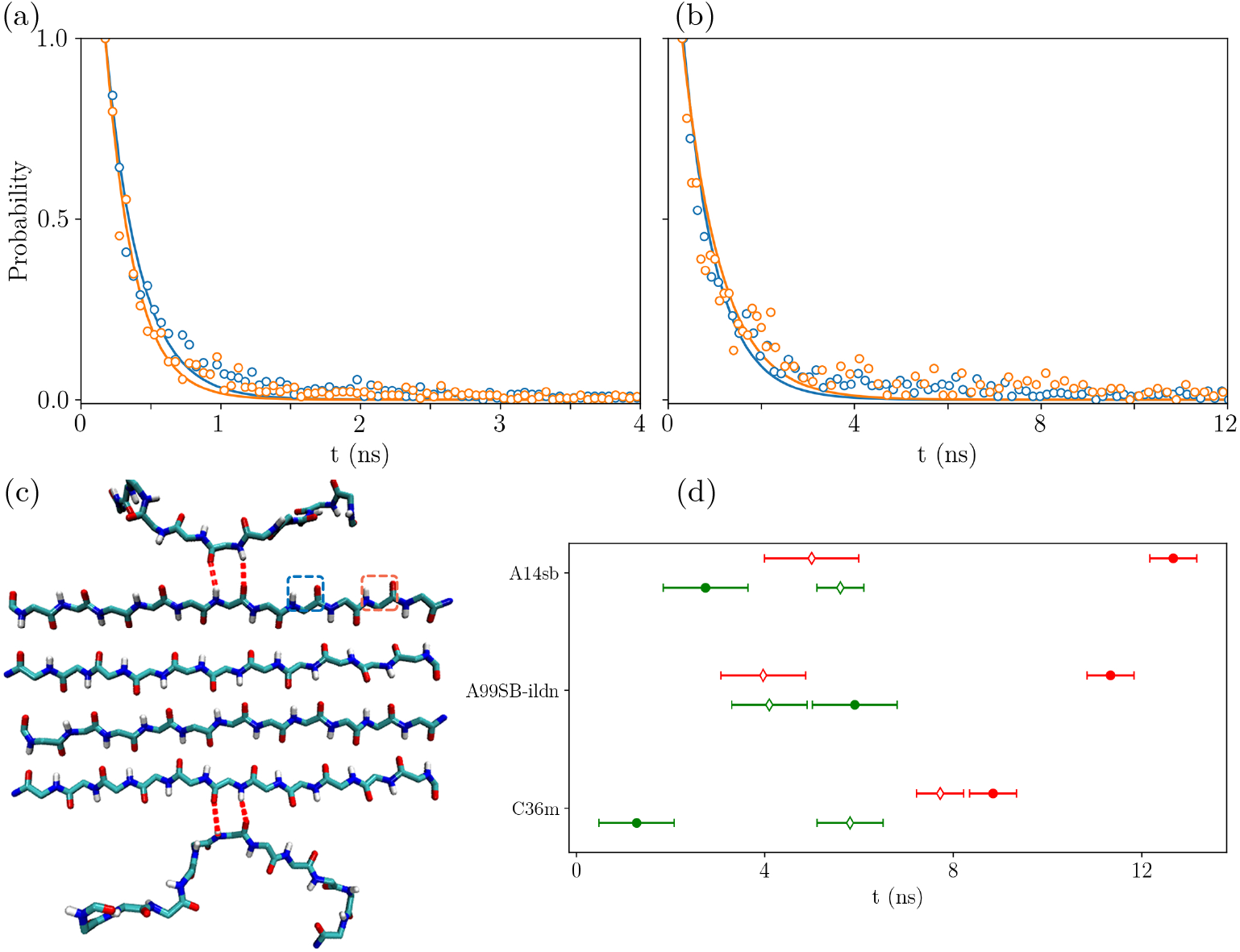
The time distributions of the unbound (a) and bound state (b) of H-bond pairs in the hexamer. Blue points indicate the H-bond pair immediately adjacent to the restrained bonds (red dashes in panel (c)) while orange points indicate the second pair from the restraints, as indicated by the by the highlighted squares in panel (c). Solid lines show exponential fits. (d) Average H-bond pair transition times associated with three different force fields. The open and closed markers represent the times of unbound and bound state of the H-bond pair respectively. Red indicates strong bonds and green indicates weak bonds.

We used three force fields to measure the kinetic rate constants. To compare force fields, it is convenient to compute the free energy of the H-bonds using the detailed balance relation

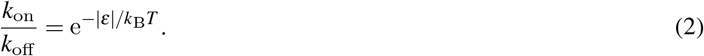

As expected, the *β* state in the dimer (Fig. 1b), representing the pre-nucleation state, is much less attractive than the post-nucleation hexamer (Fig. 1a). We refer to the bonds in the pre- and post-nucleation states as “weak” and “strong”, respectively. Following the expectations of (11) and previous theoretical analysis (37–39) we attribute the difference between these states to the entropic cost of straightening two peptide backbones to form an H-bond in the dimer compared to only one bond that must be immobilized in the post-nucleation case. This explanation is further supported by the fact that the 1.6 *k_B_T* difference between pre- and post-nucleation H-bonds in 2 out of the 3 force fields we tested agrees nearly quantitatively with the *k_B_T* ln(6) value for the backbone entropy found by fitting helix-coil transitions (40). This additional cost of forming *β*-sheet structure in the dimer will contribute to the energy barrier that must be surmounted to nucleate a fibril.

Despite the fact that each of the three force fields has been favorably evaluated for simulations of polyglutamine (41–45), they yielded widely different results (Fig. 3d, Table 1). Rather than assess the relative accuracy of these results, we use these different values as representative of sequences in different regimes of fibril stability. AMBER99SB-ILDN is the most attractive, and in fact, even the weak bonds are net attractive by *ε_w_* = −0.37*k_B_T*. When the weak bonds are attractive conformational entropy cannot contribute to the barrier. The least attractive force field is CHARMM36m, which was developed in response to overly compact ensembles in simulations of disordered proteins (25). CHARMM36m has a strong bond affinity *ε_s_* = −0.14*k_B_T*, which has insufficient attraction to observe nucleation within a single *β*-sheet and suggests that steric zipper interactions would be necessary to form stable fibrils. AMBER14SB has an intermediate affinity with weak bonds *ε_w_* = +0.71*k_B_T* that are sufficiently repulsive to create a free energy barrier, yet the strong bonds *ε_s_* = −0.93*k_B_T* provide enough attraction to stabilize a single-layered *β*-sheet.

**Table 1:** Strong and weak bond free energies, in units of *k_B_T*, calculated from kinetic parameters using Eq. (2). Strong bonds have attractive free energies ranging from −0.14 to −1.05 *k_B_T*, while weak bonds range from an attraction of −0.37 *k_B_T* to a repulsion of +1.51 *k_B_T*.

|  | C36m | A99SB-ILDN | A14SB |
| --- | --- | --- | --- |
| strong bond | $-0.14 \pm 0.01$ | $-1.05 \pm 0.18$ | $-0.93 \pm 0.16$ |
| weak bond | $1.51 \pm 0.50$ | $-0.37 \pm 0.04$ | $0.71 \pm 0.23$ |

To gain intuition into how the *β*-sheet initiation cost affects nucleation kinetics, we first construct a 2D model, consisting of a single *β*-sheet, for which we can write down the free energy and numerically simulate the kinetics. For this model we employ the parameters from the AMBER14SB force field, which are sufficiently attractive to stabilize an isolated *β* -sheet, yet repulsive enough to have a barrier.

### Model free energy shows that the number of molecules in the critical nucleus is independent of concentration

We can approximate the free energy of a single *β* sheet containing *N* = *ℓ*(*M* – 1) intermolecular H-bonds and *M* peptides as follows

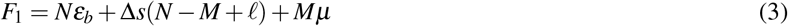

where *ℓ* is the average length of a *β*-strand, *ε_b_* is the favorable contact energy per amino acid including the H-bond and side chain interactions, Δ*s* is the conformational entropy cost to straighten a peptide unit into *β* conformation, and *μ* = *k_B_T* ln *c*/*c*_0_ is the translational entropy cost of recruiting a molecule from solution, where *c* is the peptide concentration and *c*_0_ is a reference concentration. The conformational term contains corrections due to the fact that the first H-bond on each molecule does not limit conformational degrees of freedom and the first row of *ℓ* bonds comes at a cost of 2Δ*s* per bond. This free energy can be re-written as

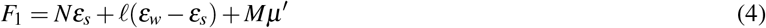

where *ε_s_* = *ε_b_* + Δ*s* is the free energy per H-bond in the interior of the *β*-sheet, *ε_w_* = *ε_b_* + 2Δ*s* is the free energy of the H-bonds in a dimer, and *μ*′ = *μ* – Δ*s*. Note that this analysis neglects minor corrections due to the staggered arrangement of H-bonds, which will be accounted for in the numerical simulation (Fig. 2). If *ε_w_* is negative (attractive), Eq. 4 is monotonically downhill for increasing *ℓ*. In this regime we recover the well-known result that one-dimensional objects lack a nucleation barrier. Therefore, we focus our attention on repulsive *ε_w_*.

The form of Eq. 4 is that of a bulk term proportional to the sheet area, *N*, competing against lines tensions that take the values (*ε_w_* – *ε_s_*)/2 = Δ*s*/2 and *μ*′/2 in the *ℓ* and *M* directions, respectively. Notably, the latter line tension is sensitive to the protein concentration. Therefore, we expect that changing the concentration will alter the shape of a nascent *β*-sheet in the same way that the surface energies of different crystal faces can result in either elongated or compact crystals.

For fixed *N*, the optimal cluster dimensions are *ℓ* = (*μ*′*N*/Δ*s*)^1/2^ and *M* = (Δ*sN*/*μ*′)^1/2^ + 1. These values can be used to find the free energy maximum

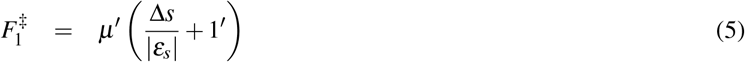

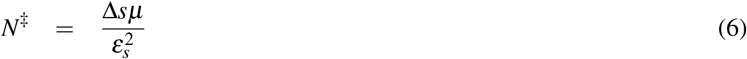

at which point the cluster has dimensions

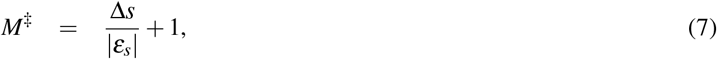

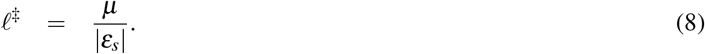

Eq. 7 has a simple interpretation if we notice that the first term is the number of internal bonds needed to pay conformational entropy cost of forming the first weak bond. Both *M*^‡^ and *ℓ*^‡^ decrease with increasing *ε_s_*, which shows that more aggregation-prone sequences have smaller critical nucleus sizes.

### Lattice model simulations show that higher concentrations result in narrower critical nuclei

Fig. 4 compares the committor, defined as the probability that a nucleus with a specified number of bonds reaches the cutoff size of 8 molecules before completely dissolving, from the kinetic model to the transition state computed from Eqs. 6 and 7. Within error the saddle point lies on the 50% committor line. Hereafter we refer to this line as the critical surface. Consistent with CNT, the critical surface shifts to smaller *N* as the concentration increases. This can also be seen in Fig. 5a, which plots the average number of H-bonds in successful trajectories crossing the critical surface. The reduction in size is not isotropic and, instead, occurs primarily in the *ℓ* direction, resulting in shorter *β*-strands as the concentration increases (Fig. 5b). This can be seen in the inset to Fig. 5b which shows that *N*^‡^/*ℓ*^‡^ is nearly constant, consistent with the prediction of Eq. 7 that *M*^‡^ is independent of concentration. Using the bond free energy from AMBER14SB gives *M*^‡^ = 2.76, which is the number of *β* strands required to overcome the initial conformational entropy cost of the dimer. From Fig. 5a, we obtain the reference concentration *c*_0_ = 44.34 *μ*M by fitting the data to *N*^‡^ in Eq. 6. This value can be used in the expression for *ℓ*^‡^ = *k_B_T*ln(*c*/*c*_0_)/(*ε_s_*), which matches the simulation results without additional parameters (Fig. 5b).

**Figure 4:**
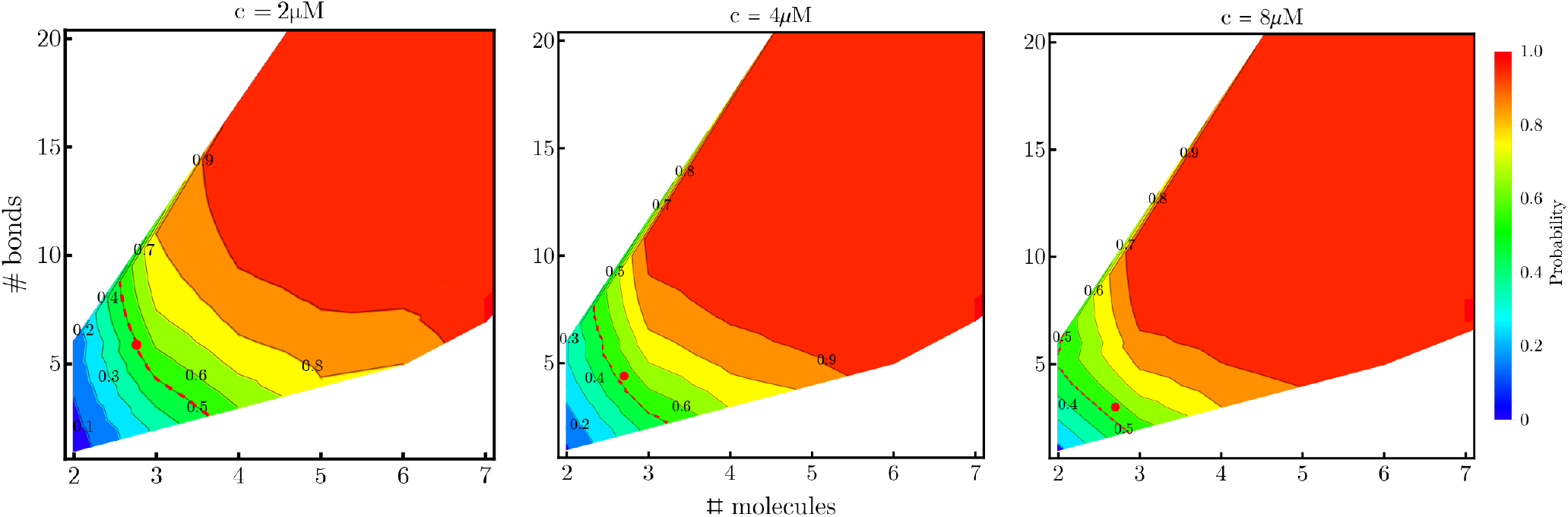
Probability of successful nucleation trajectories with parameters from the AMBER14SB force field as a function of the number of molecules and the total number of H-bonds in the cluster. Increasing concentration shifts the transition surface (50%, red dotted line) to smaller clusters and reduces the need for *β*-structure due to increased monomer deposition rate. Red dots indicate the free energy saddle points calculated from eqs. 7 and 8.

**Figure 5:**
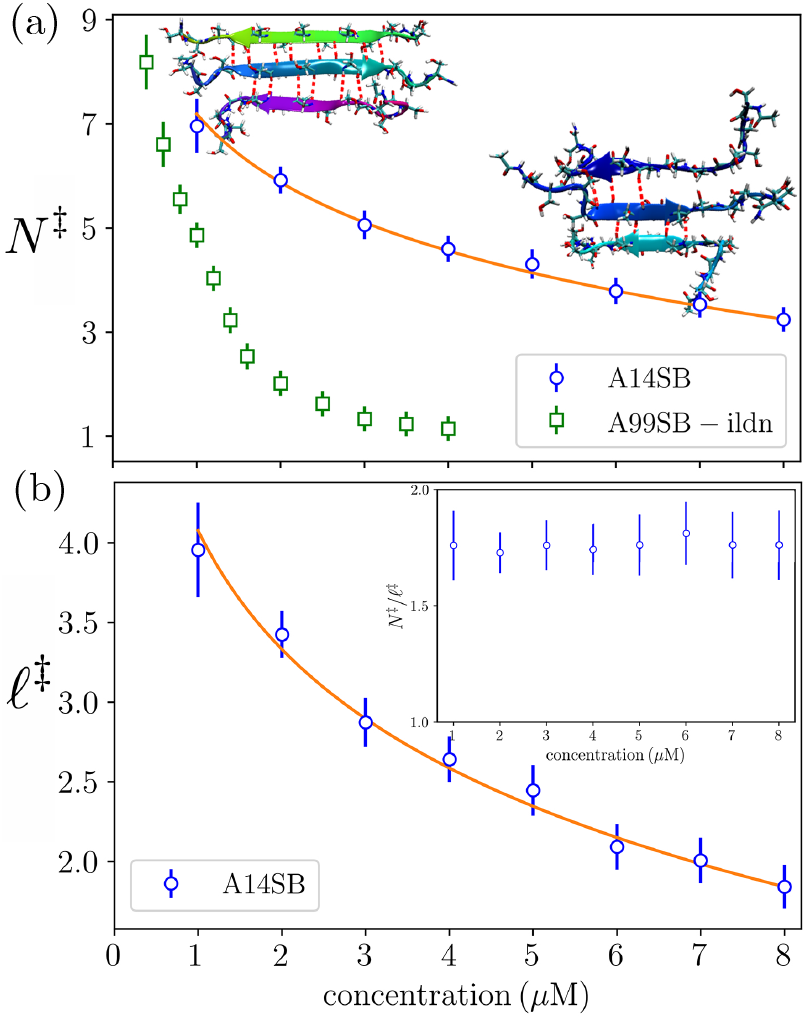
(a) Solution concentration determines the shape of the *β*-sheet core in critical nuclei as seen by the number of H-bonds in transition clusters (defined by the 50% committor). Line shows fit to Eq. 6. (inset left) Low concentration nuclei have extensive *β*-structure while higher concentration (inset right) nuclei have shorter *β*-strands. (b) The number of H-bonds per *β*-strand in the transition cluster decreases with concentration. Line shows Eq. 8. (inset) The ratio *N*^‡^/*ℓ*^‡^ is nearly independent of concentration, consistent with the prediction of Eq. 7 that the number of molecules in the critical cluster does not vary with concentration.

While *M*^‡^ is independent of concentration, it plays a crucial role in the state of the nucleus. For *M* < *M*^‡^ lateral growth of the *β* core is unfavorable, which can be seen from the derivative (*∂FF*/*∂N*)_M_ > 0. Therefore, the *β*-strands tend to remain short, which limits the ability of the nucleus to bind incoming molecules. However, for *M* > *M*^‡^, increasing *N* is favorable, so *∓* will grow by converting the disordered peptide tails to *β* conformation. The elongation of *β*-strands increases the size of the template that incoming molecules can bind to. This will increase the binding lifetime of these new molecules, which enables the transition from the nucleation stage to the elongation phase.

To understand the concentration-dependent behavior of the committor, it is instructive to examine the free energy landscape. Expressing Eq. 2 in terms of *N* and *M*, we have

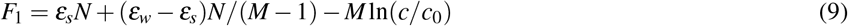

The free energy contours from this expression are plotted in Fig. 6 along with the lowest free energy path. Increasing concentration has the effect of shifting the lowest free energy pathway to smaller *N*, but does not change the number of molecules at the free energy saddle point.

**Figure 6:**
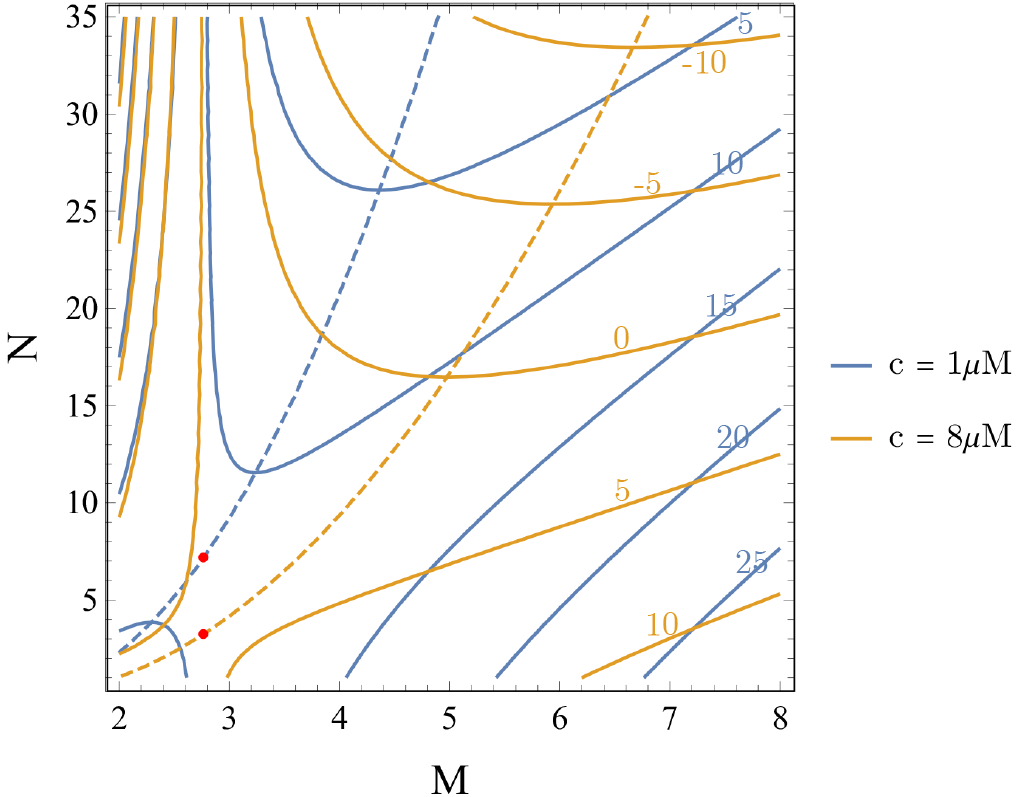
Contour plot of cluster free energy, Eq. 9 as a function of the number of molecules, *M*, and the number of intermolecular H-bonds, *N*. Increasing the concentration lowers the overall free energy, decreases the slope of the lowest free energy path (dotted lines), and shifts the free energy saddle point (red dots) to lower values of *N*.

### Multi-layered fibrils have a bigger conformational entropy penalty

The above analysis considers only a single *β*-sheet, whereas amyloid fibrils contain two or more layered sheets. To study multi-layered fibrils we use our results from the CHARMM36m force field as representative of a peptide that requires two *β*-sheets for stability. This can be seen from the single sheet parameters *ε_s_* = −0.14*k_B_T* and *ε_w_* = 1.51*k_B_T*, which would require a single sheet length *M* ⋍ 12 to overcome the initiation barrier (Eq. 7). Such a large number of molecules incurs a prohibitive translational entropy penalty unless the concentration is extremely high. Therefore, additional interactions, provided by the sidechains, are required to stabilize the fibril.

To assess the effect of the second *β*-sheet we repeat the MD simulation using a bilayer core of 8 molecules (Fig. 1c,d). Importantly, the staggered interdigitation of sidechains, where the sidechains of molecule *i* in one sheet insert between the sidechains of molecules *i* and *i* + 1 of the opposite sheet (Fig. 1), implies that molecule additions for *M* > 3 are equivalent. This uniformity allows us to apply the 2D model provided *M* ≥ 3.

From the bilayer MD simulations we find that H-bond formation events have a free energy of −1.58*k_B_T*, which suggests the following breakdown: Δ*s* = *ε_w_* – *ε_s_* = 1.65*k_B_T*, *ε_b_* = *ε_s_* – Δ*s* = −1.79*k_B_T*, and *ε_sc_* = −1.58*K_B_T* – *ε_s_* = −1.44*k_B_T* for the side chain (inter-sheet) interactions. This means that a trimer with two strands in one *β*-sheet is unfavorable by (3Δ*s* + 2*ε_sc_* + *ε_b_*)*ℓ* = 0.28*k_B_Tℓ*, whereas the addition of a fourth molecule results in a state that is favorable by (4Δ*s* + 3*ε_sc_* + 2*ε_b_*)*ℓ* = −1.3*K_B_Tℓ*. In this case initiating a new *β*-sheet with the third molecule is somewhat less favorable than a single sheet trimer, but this configuration allows for a much more favorable tetramer. Generalizing Eq. 3 we can write the free energy of the bilayer as

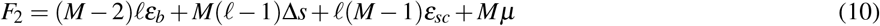

where *N* = (*M* – 2)*ℓ* is the number of intermolecular H-bonds. After rearranging, Eq. 10 takes a form similar to Eq. 4

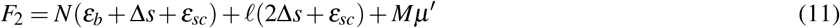

where the line tension in the *ℓ* direction has the value *γ*_2_/2 = (2Δ*s* + *ε_sc_*)/2. Therefore, a pair of layered *β*-sheets can also be described using a 2D model in which the *β*-sheet free energy has line tensions arising from conformational and translational entropy. This conclusion is valid provided the free energy maximum occurs after the initiation of the second *β*-sheet.

Both Eqs. 4 and 11 have transition state energies given by *M*^‡^*μ*′ implying that the dominant nucleation pathway is simply the one that requires the fewest number of molecules to reach the saddle point. This is given by Eq. 7 for the single sheet pathway and

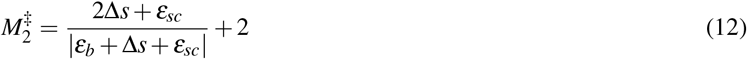

for the two-sheet pathway. Comparison of Eqs. 7 and 12 shows that the two-sheet pathway comes with a greater initial cost because there are two terminal molecules that must be straightened. However, the sidechain interaction energy somewhat reduces the initial cost below 2Δ*s* and increases the attractive energy of subsequent molecules. The nucleation barrier can be further lowered along the two-sheet pathway if the molecules are long enough to form two *β*-strands (46).

While our simulations only considered polyglutamine homopolymers, insight into the effect of amino acid sequence can be obtained from the barrier heights given by Eqs. 7 and 12. These expressions show that sidechains that stack favorably, which gives more attractive values of *ε_b_* and *ε_sc_*, will reduce the number of *β*-strands needed to overcome the entropy barrier. To a lesser extent, sidechains can also modulate Δs by perturbing the relative weights of the *β* and non-*β* regions of Ramachandran space (47).

## DISCUSSION

Experiments have shown that A*β*_16–22_ reduces the lag time of A*β*_1–40_ despite the fact that A*β*_16–22_ molecules are not incorporated in the resulting A*β*_1–40_ fibrils (48). This result is explained by our model, which shows that the *β*-sheet core contains only a portion of the aggregating molecules. This means that the nucleus will not be sensitive to the molecule length. Therefore, we predict that the lag time reduction by protein mixtures will depend on the protein concentration because at lower concentration the ordered portion of the nucleus will be larger and more sensitive to mismatches. We propose that impurity molecules incorporated during nucleation will be sites susceptible to fragmentation, enabling removal of the defects during elongation.

Our coarse-grained representation only accounts for intermolecular backbone H-bonds, however, we can speculate on the effect of molecules associating with the cluster via non-*β* contacts. Disordered binding will, in general, lower the free energy of pre-nucleation clusters thereby increasing their prevalence (49). If a *β*-sheet is to nucleate from disordered clusters (11, 12) there will be an entropy cost to orient molecules with the existing growing *β*-sheet. This rotational entropy cost will play a similar role as *μ* in Eq. 3. However, since rotational entropy does not scale with concentration, the aspect ratio of the critical *β*-sheet will be fixed if the dominant nucleation pathway goes through disordered clusters.

The large role of conformational entropy provides insights into other mechanisms for fibril nucleation, including homogenous nucleation, secondary nucleation catalyzed by fibrils, and heterogeneous nucleation at impurities or interfaces (50–54). According to CNT, binding to a surface during heterogeneous nucleation lowers the surface energy relative to homogeneous nucleation. We propose that the attachment of a peptide to a surface will lower the conformational entropy cost to form a *β*-strand. Additional evidence for this view comes from the Ramachandran plot, which is dominated by deep wells in the *α*-helix and *β*-sheet regions. Therefore, if association with a surface is favorable, the peptide will be strongly biased toward conformations compatible with *β*-sheet formation. Thus, the association of a peptide with the surface of a fibril will naturally lower the free energy of nucleation.

The nucleation mechanism in our simulations blurs the line between one-step nucleation, where condensation and ordering coincide, and two-step nucleation, where condensation precedes ordering (2, 55–57). We find *β*-sheet ordering occurs concurrently with initial cluster formation and, in fact, is necessary for molecule retention. However, the molecules remain mostly disordered. Previous simulations of fibril nucleation have shown a two-step mechanism for most cases (11). However, the model in that work represented molecules using an all-or-nothing conversion between ordered and disordered states, hence high concentration or non-specific attractions were necessary to surmount the conversion barrier. Our work shows that a partial conversion to the *β*-state is an important mechanism to lower the nucleation barrier, so nuclei are likely to have both ordered and disordered regions at all stages.

In conclusion, we have shown that the nucleation of *β*-sheets, an essential step in the formation of amyloid fibrils, is limited by both conformational and translational free entropy. These factors can be described by classical nucleation theory provided the *β*-sheet is viewed as a cluster that can grow either by the addition of molecules along the fibril axis or by the conversion of peptide units to the *β*-state in the lateral direction. We hope that this framework can be used to gain intuition about fibril nucleation in a variety of contexts.

## Author Contributions

T.M.P. and J.D.S. designed the study and wrote the paper. T.M.P. performed the research.

## ACKNOWLEDGMENTS

This work was supported by National Institutes of Health Grants R01GM107487 and R01GM141235. J.D.S. acknowledges valuable discussions with Randal Halfmann and Martin Muschol. T.M.P. would like to thank Zhiguang Jia for fruitful discussions. The simulations for this project were performed on the Beocat Research Cluster at Kansas State University, which is funded in part by NSF Grants CHE-1726332, CNS-1006860, EPS-1006860, and EPS-0919443.

